# VAMP8 Deficiency Attenuates AngII-Induced Abdominal Aortic Aneurysm Formation via Platelet Reprogramming and Enhanced ECM Stability

**DOI:** 10.1101/2025.02.03.635525

**Authors:** Shayan Mohammadmoradi, Elizabeth R. Driehaus, Hammodah R. Alfar, Smita Joshi, Sidney W. Whiteheart

## Abstract

**BACKGROUND:** As vascular sentries, platelets, and their ability to release a host of bioactive molecules, are critical for vascular homeostasis as well as hemostasis. Despite data linking platelet activation to abdominal aortic aneurysms (AAA) and rupture, the underlying mechanisms remain poorly understood. This study addresses the hypothesis that VAMP8, the primary v-SNARE controlling platelet exocytosis, contributes to AAA formation.

**METHODS AND RESULTS:** In an AngII-infused hypercholesterolemic mouse model, we observed significant platelet consumption, indicated by decreased platelet counts at both acute (5-day) and chronic (28-day) time points. Platelets accumulated at sites of elastin degradation and within false lumens of the abdominal aorta after 28 days of AngII infusion. Bulk RNA sequencing analysis of washed platelets and their releasates after 5 days of AngII infusion revealed significant transcriptomic changes, suggesting rapid reprogramming of platelet function. Parallel RNA-seq analysis of suprarenal aortic tissue highlighted changes in genes associated with extracellular matrix (ECM) organization, inflammation, and platelet signaling, linking platelets to vascular remodeling suggesting a “platelet-aorta axis”. Laser speckle imaging in a FeCl₃ injury model confirmed that VAMP8 deficiency impaired platelet function, resulting in delayed thrombosis. *In vivo* experiments demonstrated that VAMP8⁻/⁻ mice were protected against AngII-induced AAA and aortic rupture. Aortic diameter analysis further revealed that VAMP8 deficiency significantly attenuated AngII-driven aortic pathology. RNA-seq analysis of platelets and aortic tissue suggests that loss of VAMP8 affects expression of genes controlling ECM degradation and aortic wall stability consistent with the protective effect of VAMP8 loss on AAA.

**CONCLUSION:** Short-term AngII infusion appears to reprogram the platelet transcriptome, which may affect the aorta and contribute to AAA formation. Controlling cargo release from platelets via VAMP8 deficiency results in profound attenuation of aortic aneurysms. This introduces a novel paradigm for understanding the impact of reprogrammed platelet cargo secretion and function in aortopathies.

**Graphical Abstract:** 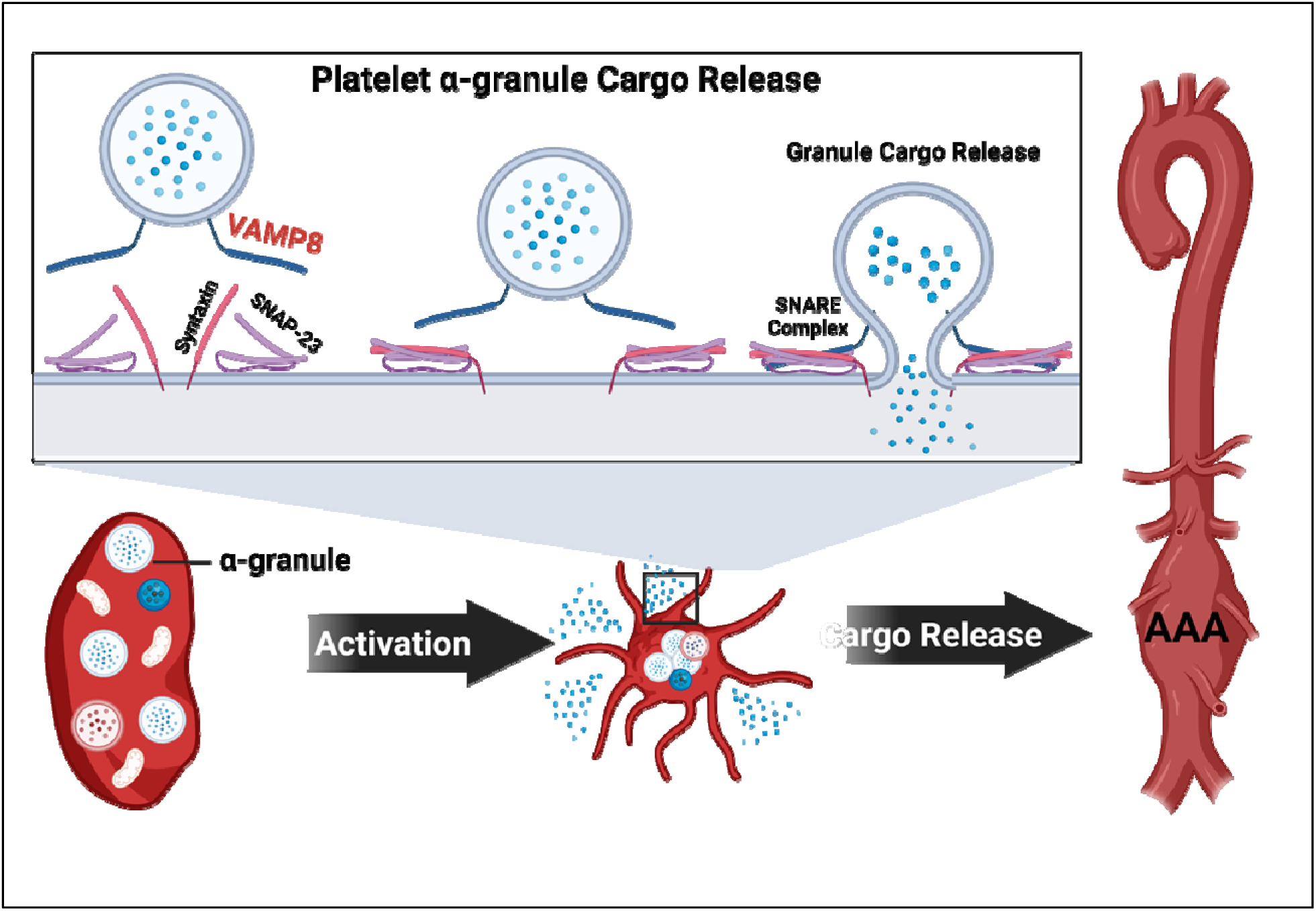

**Highlights:**

- Platelet transcriptome is altered at early aneurysmal stage.
- VAMP8 deficiency attenuates aortic aneurysms, potentially via enhanced ECM stability.
- VAMP8 deficiency significantly alters various genes contributing to aortic wall structure and stability in both platelets and suprarenal aortic tissue.

## INTRODUCTION

Increased platelet activation is a characteristic feature of various cardiovascular complications, including abdominal aortic aneurysm (AAA). While platelets are traditionally known for their roles in thrombosis and hemostasis, their involvement in aortic aneurysms has only recently been recognized. ^1–6^ Several studies propose that antiplatelet therapy might decelerate the progression of established AAA ^1,3,4,6,7^, and preclinical models have observed a reduced rupture rate following antiplatelet treatment in angiotensin II (AngII)-induced AAA. ^3,4,8^ Although the precise pathways of AAA initiation remain elusive, early events likely involve a vascular injury that results in platelets recruitment at the site of injury, initiates localized inflammatory responses, attracting macrophages and leads to progressive medial layer deterioration. ^9,10^ Further, a prominent feature of AAA is the accumulation of an intraluminal thrombus (ILT) along the luminal surface ^11^. The aggregation of inflammatory cells and anucleate cells such as platelets within the ILT can compromise the vessel wall, heightening the risk of aortic rupture. ^11–14^ Moreover, disturbed blood flow within the aneurysmal aorta can further trigger platelet activation. ^2^ Despite existing reports from preclinical animal models and small clinical trials on the involvement of platelets and platelet inhibition in AAA, there has been minimal focus on understanding how platelets contribute to AAA development, particularly their initiating role in vessel wall damage through the release of granule cargo after activation.

Many platelet functions in thrombosis and hemostasis are mediated by their releasate, generated by the exocytosis of cargo from three types of granules: dense granules, α-granules, and lysosomal granules. ^15,16^ Controlling the release of this cargo holds promising therapeutic implications for treating various platelet-related disorders. The exocytosis of granules is mediated by a group of membrane proteins known as soluble N-ethylmaleimide-sensitive factor attachment protein receptors (SNAREs), which facilitate the fusion of granule membranes with the plasma membrane. ^15,17–19^ Our group and others have identified vesicle-associated membrane protein 8 (VAMP8) as the primary VAMP responsible for platelet granule secretion. ^18,20–22^ Notably, platelets lacking VAMP8 exhibit substantial deficiencies in α-granule and lysosome release, with variable effects on thrombosis and hemostasis. ^18,22^

This study investigates the role of platelet granule cargo release in the development of AngII-induced AAA. We found that short-term AngII infusion triggers platelet reprogramming toward a hyperactive state, reflected in parallel transcriptomic changes within abdominal aortic tissue. Notably, we demonstrated that genetic deletion of VAMP8 prevents AngII-induced AAA formation, potentially by downregulating platelet pathways critical to aortic wall integrity. These results underscore that platelet activation is essential not only for AAA formation but also for maintaining the structural stability of the aortic wall.

## METHODS

### Data Availability

Comprehensive descriptions of the experimental procedures are provided in the Supplemental Methods section. Numerical data supporting the findings of this study are available in the Supplemental Excel Files. All raw datasets and analytical methods can be obtained from the corresponding author upon reasonable request.

### Animal Models and Treatment

VAMP8-deficient (VAMP8^-/-)^ mice (in-house), along with C57BL/6J (Jackson Lab #000664) or appropriate age-matched littermate wild-type (WT) controls, were used in this study. The VAMP8 genotype of each mouse was determined by PCR using DNA extracted from the tail tip, as previously described. ^22^ Mouse housing conditions are described in Supplemental Table II. Briefly, study mice were housed in individually ventilated cages (5 mice per cage) on a 14:10 hour light cycle. Teklad Sani-Chip (catalog No. 7090A; Inotiv) was used as cage bedding. Mice were fed a normal rodent laboratory diet (Diet #2918, Harlan Teklad) and provided with drinking water from a reverse osmosis system ad libitum unless otherwise noted. Detailed mouse information and inclusion and exclusion criteria are provided in the Major Resources Tables in the Supplemental Material. Due to the low incidence of experimental AAA in female mice within the AngII-induced pathology model, all studies were conducted on male mice. ^23^ All mouse experiments were conducted in accordance with the Animal Research: Reporting of In Vivo Experiments (ARRIVE) guidelines and were approved by the University of Kentucky Institutional Animal Care and Use Committee.

### Osmotic Mini Pump Implantation and Angiotensin II Infusion

Male mice aged 8 to 12 weeks were randomly assigned to receive either saline or angiotensin II (AngII, 1,000 ng/kg/min; H-1705, Bachem) via continuous infusion using a subcutaneously implanted osmotic pump. An Alzet osmotic mini-pump (Model 2004) was implanted subcutaneously into the interscapular space for this purpose. ^24^ Surgical staples used to close the incision sites were removed 7 days after surgery. Postoperative pain was managed by the application of topical lidocaine cream (4% wt/wt; LMX4, 0496-0882-15, Eloquest Healthcare, Inc). At the end of the infusion period, mice were anesthetized and euthanized for the collection of blood and organs. All study mice were checked at least once daily. Necropsies were performed immediately to determine the cause of death if a carcass was found. Aortic rupture was defined as the presence of extravascular blood accumulated in a body cavity. The location of blood egress was determined by the location of the blood clot and a discernible disruption of the aortic wall.

### Hypercholesterolemia Induction by Injection of Adeno-associated Viral Vector

To induce hypercholesterolemia for the AAA studies, mice aged 6 weeks were administered an adeno-associated viral (AAV) vector expressing a gain-of-function mutation (D377Y) in mouse PCSK9 (equivalent to human PCSK9D374Y gain of function mutation) and were fed a Western diet (WD) containing saturated fat (milk fat 21% wt/wt) and cholesterol (0.2% wt/wt; Diet # TD.88137, Inotiv). University of Pennsylvania Viral Vector Core produced the AAV vectors (serotype 8). Two weeks post-PCSK9 AAV infection, mice were infused with AngII (1,000 ng/kg/min) or saline for 28 days using subcutaneously implanted Alzet mini-osmotic pumps. ^25,26^

### FeCl_3_ Carotid Injury Model and Laser Speckle Imaging

This FeCl_3_-induced carotid injury model was performed as described ^27^. Mice (8–12 weeks old) were anesthetized using Avertin (tribromoethanol; 0.2 g/kg, i.p.). The left carotid artery was exposed using blunt dissection under a dissecting microscope and a thin strip of translucent plastic was placed underneath the artery to isolate the vessel from the surrounding tissue. The entire surgical stage was then positioned underneath a laser speckle perfusion imager (PeriCam PSI HR, Perimed, Järfälla, Sweden) to monitor blood flow. After reading distance was set, the surgical field was dried to prevent dispersion of the FeCl_3_ solution from filter paper to neighboring tissue. Thrombus formation was induced by placing a round pad of filter paper (1 mm diameter) saturated with 6% FeCl_3_ solution (1 μL) on the top of the vessel for 3 min. After 3 min, the paper was removed, and normal saline was added to dilute any residual FeCl_3_ and facilitate blood flow. The recording was stopped 25 min after the removal of the FeCl_3_ pad. Perfusion was measured from a 0.2 cm^2^ ROI positioned over the injured area and normalized to an identically-sized background ROI in each frame. Flow was expressed as a percentage of the maximum perfusion value for each recording, and time to occlusion and area under the curve were used as metrics of thrombus formation and relative stability.

### Platelet Preparation from Mouse Blood

Mice were euthanized by CO_2_ inhalation. The thoracic region was then exposed, and blood was collected from the right ventricle via a syringe with approximately 120 μL of 3.8% sodium citrate with apyrase (0.2 U/mL) and prostaglandin I_2_ (PGI_2_; 2 μg/mL). Platelet concentration was calculated using a Z2 Counter (Beckham Coulter, Inc. Brea, California). Whole blood counts were also performed using IDEXX ProCyte DX Hematology Analyzer.

### RNA Isolation and Sequencing

Washed mouse platelets isolated from WT or VAMP8^-/-^ mice were subjected to RNA isolation using the mirVana™ Total RNA Isolation Kit, followed by bulk RNA sequencing analysis. Three sets of washed platelets were pooled into one sample for RNA sequencing, with two total samples used for each group. For aortic samples, total RNA was extracted from the suprarenal aortas of VAMP8^-/-^ and WT mice after 5 days of AngII infusion. Aortas were perfused with saline, periaortic tissues were removed, and aortas displaying intramural hemorrhage were excluded. Aortic samples not displaying overt pathologies were used for transcriptomic analysis. Four aortic samples per group were incubated with RNAlater solution (No. AM7020; Invitrogen) for 1 hour. Subsequently, mRNA was extracted using RNeasy Fibrous Tissue Mini Kits (No. 74704; Qiagen) and shipped to Novogene (CA) for mRNA sequencing. Messenger RNA was purified from total RNA using poly-T oligo-attached magnetic beads. After fragmentation, the first strand cDNA was synthesized using random hexamer primers, followed by the second strand cDNA synthesis using either dUTP for directional library or dTTP for non-directional library. cDNA libraries were sequenced by NovaSeq X Plus (Illumina). The human platelet RNA sequencing data for control subjects (n = 7) and AAA patients (n = 6) was obtained from a previous publication. ^2^

### Statistical Analysis

Normality and homogeneity of variance were evaluated using the Shapiro-Wilk and Brown-Forsythe tests, respectively. For comparisons of means between two groups or multiple groups, a two-sided Student’s t-test or a two-way analysis of variance (ANOVA) with the Holm-Sidak multiple comparison test was employed, respectively. Welch’s t-test was used for data that passed normality test, but failed to satisfy equal variance assumption to compare two-group means. For data that did not pass either normality or equal variance test, we applied Mann-Whitney U test for two-group comparisons or Kruskal-Wallis one-way ANOVA followed by Dunn’s method for multiple-group comparisons. For aortic rupture-based mortality between groups the comparison was made using the log-rank test. Statistical analyses were conducted using SigmaPlot version 15.0 (SYSTAT Software Inc.). The number of mice per experiment is detailed in each figure legend. Data are presented as mean ± SEM. A p-value of less than 0.05 was considered statistically significant. Mice were randomly assigned to study groups. An independent investigator, blinded to the study group information, verified all experimental data. Control data were acquired concurrently with experimental data for statistical comparisons.

For RNA Sequencing analysis, differential expression analysis was performed using the DESeq2Rpackage (1.20.0). The resulting P-values were adjusted using the Benjamini and Hochberg’s approach for controlling the false discovery rate. Genes with an adjusted P-value <=0.05 found by DESeq2 were assigned as differentially expressed. (For edgeR without biological replicates) Prior to differential gene expression analysis, for each sequenced library, the read counts were adjusted by edgeR program package through one scaling normalized factor. Differential expression analysis of two conditions was performed using the edgeR R package (version 3.22.5). The P values were adjusted using the Benjamini & Hochberg method. Corrected P-value of 0.05 and absolute foldchange of 2 were set as the threshold for significantly differential expression. Gene Ontology (GO) enrichment analysis of differentially expressed genes was implemented by the clusterProfiler R package, in which gene length bias was corrected. GO terms with corrected P value less than 0.05 were considered significantly enriched by differential expressed genes. We used clusterProfiler R package to test the statistical enrichment of differential expression genes in KEGG pathways. Pathways enriched among differentially expressed genes were visualized as bubble plots, with enrichment scores or p-values displayed on the x-axis, pathway names on the y-axis, bubble size corresponding to the gene count, and color indicating statistical significance. Visualization of RNA sequencing data were done using Python (version 3.8 or higher) libraries including *matplotlib*, *seaborn*, and *plotly*.

Additional methods are in the Supplemental Material.

## RESULTS

### Increased Platelet Consumption and Thrombosis in AngII-Induced AAA Mouse Model

To investigate the effect of AngII infusion on platelets, we asked whether increased platelet activation and potential consumption could be observed under short-term (5-day) and full pathological (28-day) conditions. At both time intervals, we observed reduced platelet counts, along with a significantly higher mean platelet volume (MPV) in the aneurysmal condition **(Figure 1 A and B)**. Such data could indicate enhanced platelet consumption and/or sequestration.

**Figure 1:**
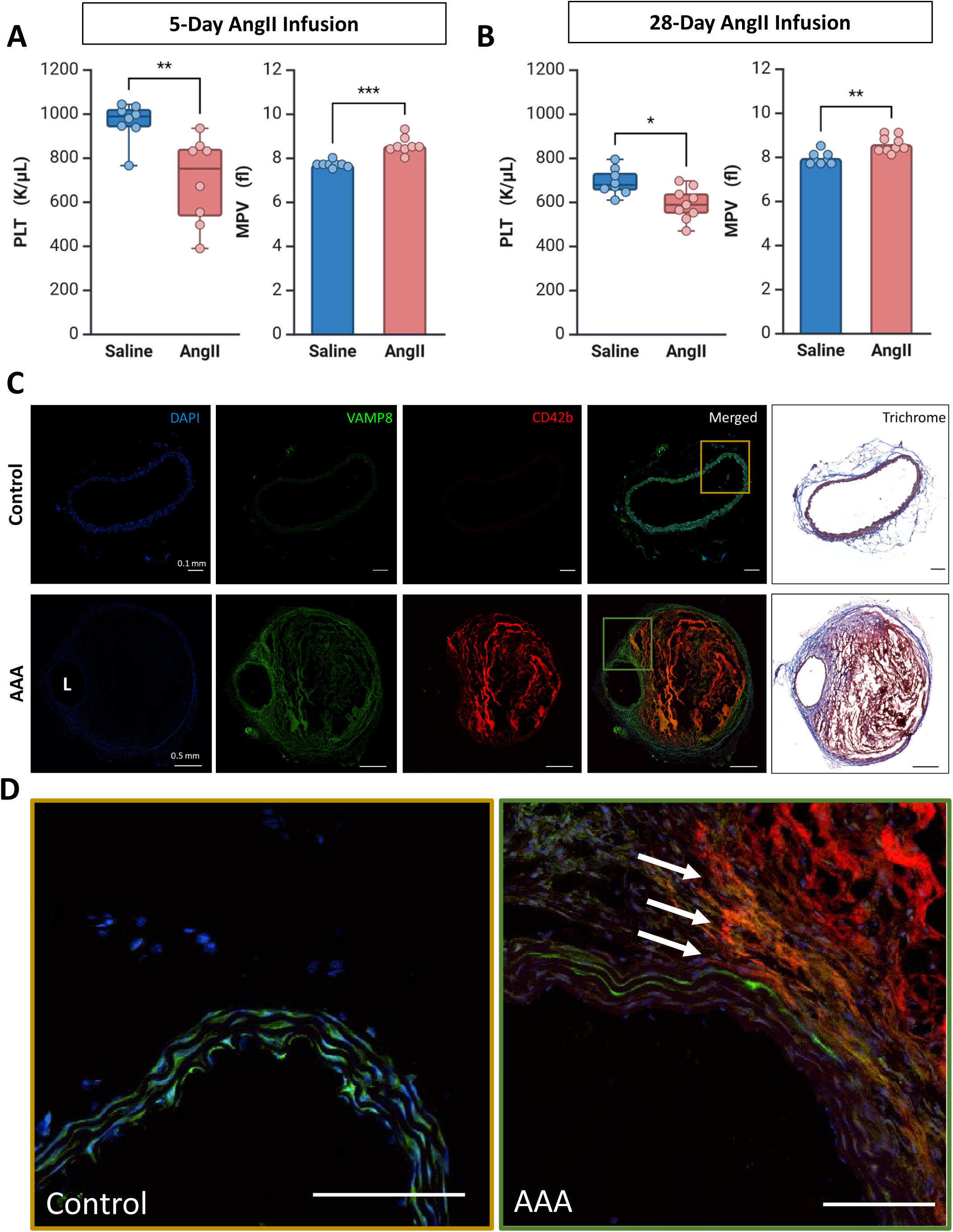
Platelets Distribution in AngII-induced Aneurysmal Abdominal Aorta. Platelet count and mean platelet volume (MPV) was determined by complete blood count from male mice in 5 **(A)** or 28 **(B)** days of angiotensin II (AngII, 1000 ng/kg/min) infusion (n=8-9/group). **(C)** Representative maximum-intensity projection images from immunofluorescence and trichrome histological analyses of suprarenal aortic sections from 28-day saline or AngII-infused mice. Histochemistry and fluorescence sections represent varying depths within the identical specimen (n=3/group). **(D)** Zoomed images highlight areas of interest, indicating potential platelet migration into the medial layer in the AAA group. Data are presented with circles and box plots as mean ± SEM. P-value was determined by Welch’s t-test. ** P ≤ 0.01, *** P ≤ 0.001. Arrows indicate areas with platelet infiltration. L: Lumen, AAA: abdominal aortic aneurysm

### Platelet Accumulation at Sites of Aortic Wall Disruption in AngII-Induced AAA

We next examined the distribution of platelets in suprarenal aortic tissue with aneurysm. Suprarenal aortas from mice infused with AngII (1,000 ng/kg/min) for 28 days were immunostained with anti-CD42b (a platelet marker) and anti-VAMP8 (primary platelet v-SNARE) antibodies. Consecutive sections were stained with Masson’s trichrome. In saline-infused mice, platelets and VAMP8 were nearly absent from the healthy abdominal aorta. After 28 days of AngII infusion, there was a notable increase in the platelets and VAMP8 abundance in the aneurysmal tissue, especially in the thrombus region. Platelets were significantly concentrated at sites of elastin breaks and false lumen formation **(Figure 1C).** Additionally, platelets were observed in the medial layer of disrupted aortic tissue, suggesting migration into the aortic elastin layers, particularly at sites of elastin breaks and false lumen formation **(Figure 1D).** These findings further indicate platelet activation and recruitment to the aortic wall in mice with AngII-induced AAA.

### Platelet Transcriptome Is Altered Toward a Hyperactive State in Early and Late AAA Pathological Conditions

Given the decrease in platelets and its accumulation in the aneurysmal sac, we next determined how platelets change during aneurysm initiation. We isolated platelets from age-matched WT mice infused with AngII or saline for 5 days and performed Bulk RNA sequencing (n=4 per group; each sample consisted of pooled platelets from 2 mice). Short-term AngII infusion resulted in significant transcriptomic changes in platelets. We identified a total of 997 differentially expressed genes (DEGs) in response to AngII, with 550 genes upregulated and 447 downregulated **(Figure 2A)**. Reactome pathway analysis of the DEGs revealed significant enrichment in pathways related to “hemostasis,” “platelet activation,” and “platelet degranulation” **(Figure 2B).** Further analysis of a curated list of genes encoding platelet granule cargo showed significant fold-change increases, indicating a strong effect on platelet activation and function **(Figure 2C)**. These results suggest that short-term AngII infusion (5 days) leads to alterations that reflect platelet reprogramming. Reanalysis of previously published RNA-seq data of human platelets^2^ from healthy individuals (n=7) and patients with AAA (n=6) revealed similar enrichment of Reactome pathways such as “platelet activation” and “degranulation” **(Figure 2D)**. This was accompanied by significant upregulation of genes relevant to platelet activation and function, aligning well with our animal studies **(Figure 2E)**. The concordance between the human and mouse data suggests that platelet hyperactivity is a conserved feature in AAA pathology.

**Figure 2:**
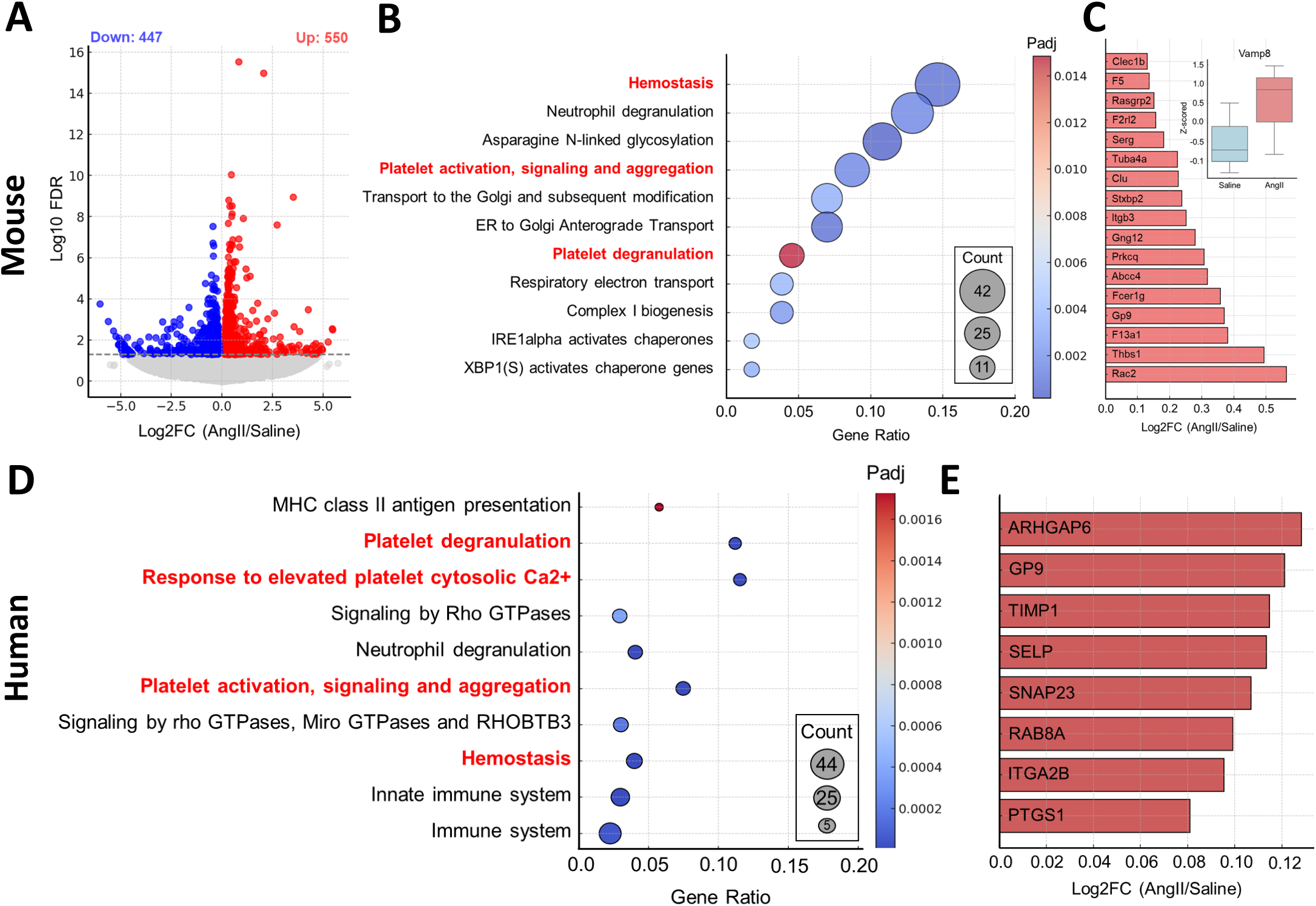
Platelet Transcriptome Is Altered Toward a Hyperactive State in Early and Late AAA Pathological Conditions. **(A)** Volcano plot for differentially expressed genes between washed platelets of 5-day saline- and angiotensin II (AngII)–infused mice (n=4/group – each sample is consisted of pooled platelets from 2 mice). **(B)** The 10 most significant Reactome terms. **(C)** Log fold change of genes related to platelet function compared between saline and AngII infusion **(D)** The 10 most significant Reactome terms of human platelet RNA sequencing extracted from healthy individuals (n = 7) and patients with AAA (n= 6). **(E)** Log fold change of genes related to platelet function compared between platelet transcriptome of healthy individuals vs. AAA patients.

### AngII-Induced Transcriptomic Changes in Aortic Tissue Reveal a Platelet-Aorta Axis

To further uncover the molecular mechanisms of AngII-induced aortic pathologies and its relevance to a “platelet-aorta axis,” we performed bulk RNA sequencing of the suprarenal aortic segments—the area prone to AngII-induced AAA formation. Suprarenal aortic tissues were harvested at day 5 of either saline or AngII infusion after perfusion, excluding samples showing intramural hemorrhage or dissection. Compared with saline infusion, AngII infusion altered the expression of 2,117 genes, with 1,208 upregulated and 909 downregulated **(Figure 3A)**. As expected, Reactome pathway analysis and Gene Ontology (GO) enrichment analyses of the DEGs indicated significant enrichments in terms related to extracellular matrix (ECM) organization, degradation of ECM, cytokine binding, and receptor-ligand activity, **(Figure 3 B, C).** We next examined ECM-related molecules, chemokines, cytokines, and genes relevant to platelet function—including *Spp1, Gp9, Serpine1, Plau, Thsp1, and Vamp8*—all with clear relationships to platelets **(Figure 3D)**. These genes showed significant fold-change increases in response to AngII. To further explore the similarities between the transcriptomic alterations in the “platelet-aorta axis,” we analyzed the DEGs from both platelet and aortic tissues, identifying 58 genes that were commonly upregulated **(Figure 3 E, G)**. GO molecular pathway analysis of these commonly upregulated genes indicated enrichment in terms strongly associated with platelets and their function **(Figure 3F)**, such as “platelet formation,” “wound healing,” and “inflammatory response.” These findings suggest a coordinated response between platelets and the aortic tissue in AngII-induced AAA, highlighting the potential role of platelet-related cargo in the pathogenesis of aortic aneurysms.

**Figure 3:**
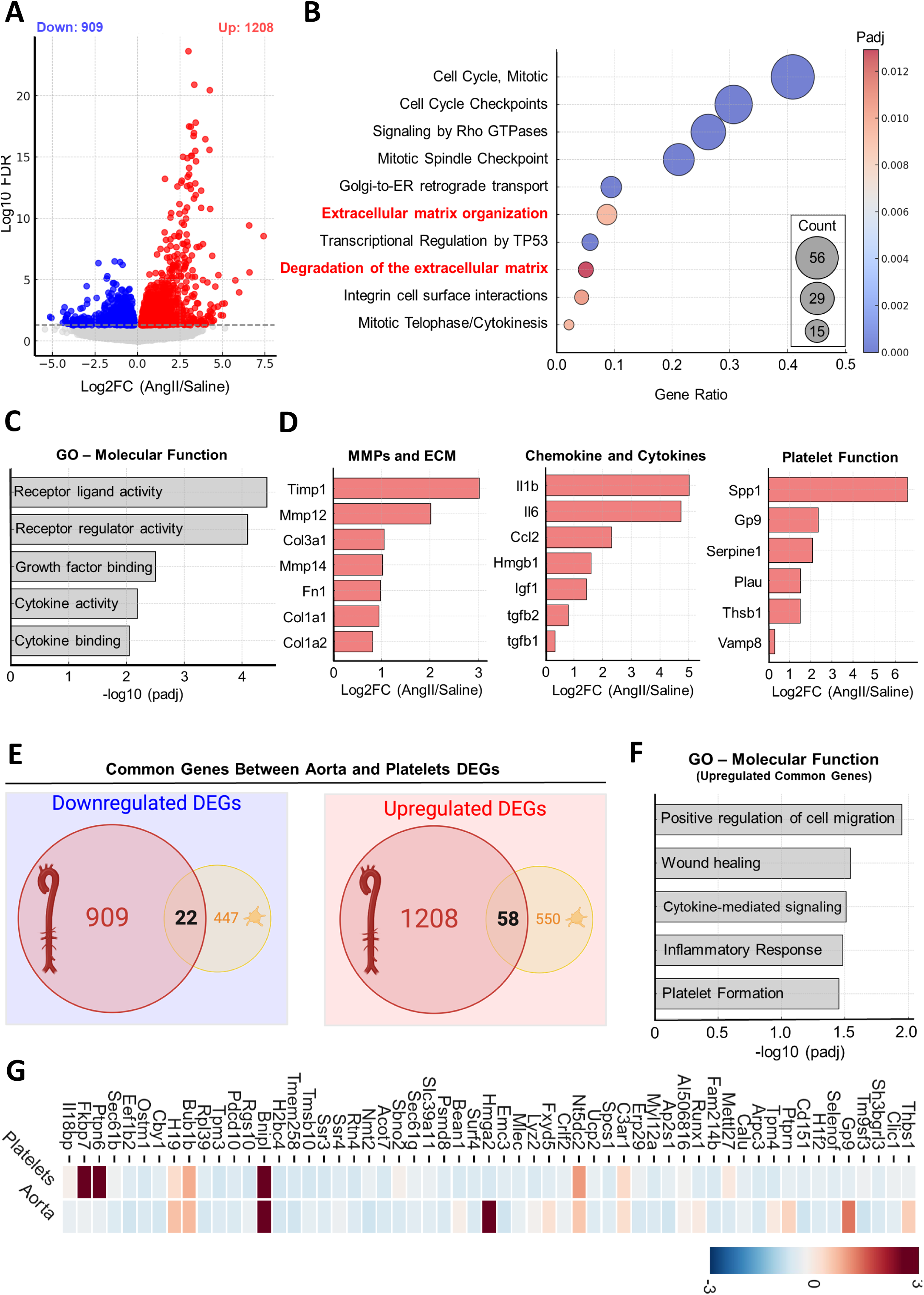
AngII-Induced Transcriptomic Changes in Aortic Tissue Reveal a Platelet-Aorta Axis. **(A)** Volcano plot for differentially expressed genes between suprarenal aortas of saline- and angiotensin II (AngII)–infused mice (n=4/group). **(B)** The 10 most significant Reactome terms. **(C)** Top 5 annotations of enrichment analyses in the Gene Ontology molecular function. **(D)** Log fold change of genes related to matrix metalloproteinase (MMP) and extracellular matrix (ECM), chemokines and cytokines, and platelet function compared between saline and AngII infusion. **(E)** *Venn diagrams* for shared differentially expressed genes between downregulated and upregulated aortic and platelet datasets. **(F)** Top annotations of Gene Ontology molecular function enrichment analysis in the common upregulated differentially expressed genes of aorta and platelets. **(G)** Heatmap with Z score coloring for common upregulated genes between platelet and aortic samples.

### VAMP8 Deficiency Delays Occlusive Thrombosis without Impairing Hemostasis

Previous studies showed that VAMP8 is the primary v-SNARE for platelet exocytosis and its loss significantly affects α-granule release more than dense granule release. ^22^ Consistently, this defect has minimal effect on bleeding in some models but has been reported to delay occlusive thrombosis in arterial model injury. ^18,28^ To better define this, we reexamined occlusive thrombosis using an imaging method that focuses on the particles flow; laser speckle imaging system. We performed the FeCl₃-induced (6%) carotid artery injury model as previously described. ^18,29^ Our results show that in the FeCl₃-induced carotid occlusion model, WT mice formed fully occlusive thrombi rapidly, while VAMP8-null mice exhibited delay, with occlusion occurring at approximately 20 min **(Figure 4 A-C).** Notably, occlusion times in the VAMP8^⁻/⁻^ mice were much more heterogeneous, suggesting an effect on clot stability and growth. These findings indicate that while VAMP8 deficiency delays thrombus formation in large vessels, it does not impair overall hemostasis, as clotting still occurs, albeit more slowly. Thus, VAMP8 deficient platelets are able to contribute to normal responses to vascular damage but at a reduced efficiency, which is sufficient for hemostasis, although with delayed kinetics.

**Figure 4:**
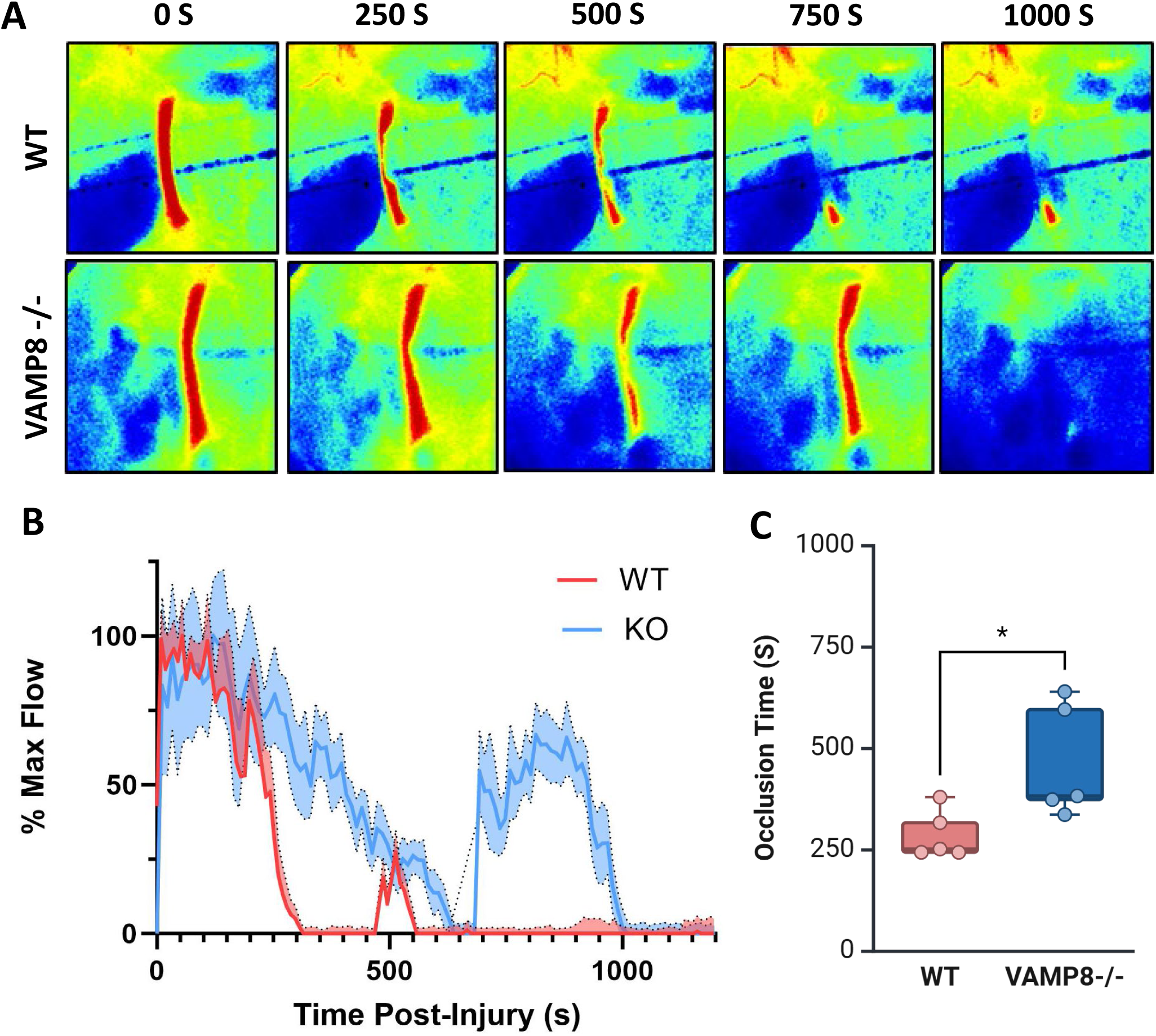
Depleting VAMP8 Affects Vascular Homeostasis. Thrombus formation in the carotid artery was induced by topical application of 6% FeCl3 and blood flow was monitored in WT (n=5) and VAMP8^⁻/⁻^ (n=5) mice for 25 mins. **(A)** Representative laser speckle perfusion images of WT and VAMP8-/- carotid arteries at t = 0, 5, 15, and 25 mins post-injury. **(B)** Representative flow traces of WT and VAMP8^-/-^ carotid arteries over a period of 25 mins post-injury. **(C)** Time to occlusion is shown and P value (log-rank test) is indicated. Data including flow traces are expressed as mean ± SEM. * P ≤ 0.05

Given the impact of VAMP8 deficiency on occlusive thrombosis, we investigated its effect on the early stages of atherosclerosis development. Animals were injected with AAV-PCSK9 (D377Y) and fed a Western diet for 4 weeks. VAMP8-deficient male mice demonstrated a significant reduction in early-stage atherosclerosis progression (Supplementary Figure 2 A-C). These findings suggest that the delayed thrombus formation and reduced clot stability in VAMP8-deficient mice may contribute to decreased arterial inflammation and lesion formation, thereby attenuating early atherosclerosis progression.

### VAMP8 Deficiency Profoundly Protects Against AngII-Induced AAA

Given the significant role of VAMP8 in platelet cargo release and its sequela, we sought to define the role of VAMP8 and defective platelet cargo release in AAA pathophysiology. We induced AAA using the AAV-PCSK9(D377Y)/AngII model in WT and VAMP8^⁻/⁻^ mice. They were fed a high-fat diet, and infused with AngII (1,000 ng/kg/min) or saline two weeks post-PCSK9 AAV infection **(Figure 5A).** The VAMP8^⁻/⁻^ mice showed similar changes in systolic blood pressure **(Supplementary Figure 1A)** and cholesterol levels in the plasma **(Supplementary Figure 1B)** were increased similarly to WT, demonstrating that loss of VAMP8 is not overtly change the systemic pathologies induced by the model. After 28 days, VAMP8^⁻/⁻^ mice were fully protected against AngII-driven aortic rupture **(Figure 4B)**. VAMP8 deficiency profoundly attenuated AngII-induced aortic dilation in the thoracic and abdominal aorta, as demonstrated by ultrasound imaging **(Figures 4C and 4D).** *Ex vivo* analysis of the aorta validated that VAMP8 deficiency (n=15; aortic diameter: 0.9 mm ± 0.03) significantly attenuated AngII-induced AAA compared to control mice (n=9; aortic diameter: 2.0 mm ± 0.2) **(Figures 5E and 5F).** These studies demonstrate that loss of VAMP8 attenuates aortic aneurysm formation.

**Figure 5:**
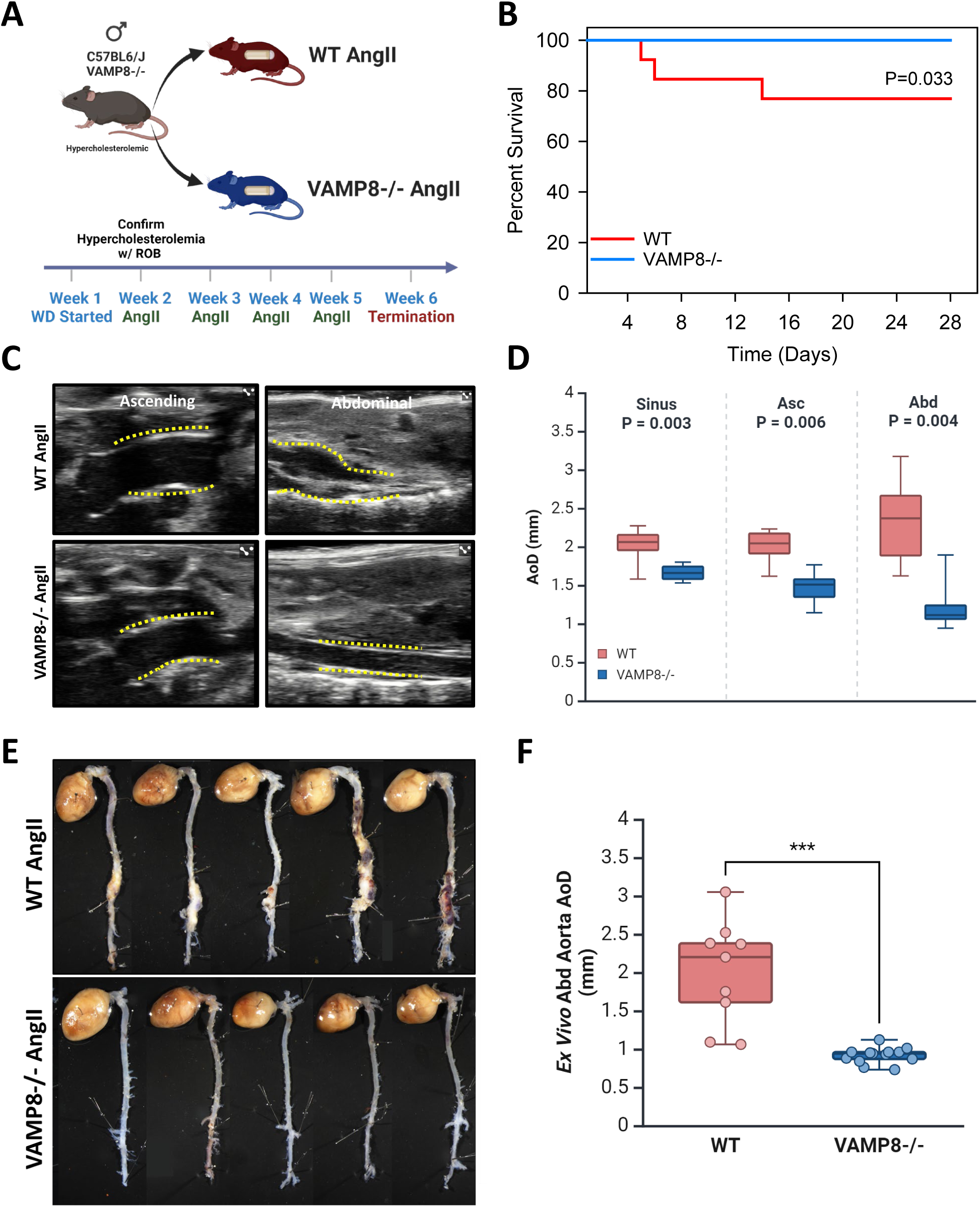
VAMP8 Deficiency Profoundly Attenuated AngII-induced Aortic Rupture and Aortic Aneurysms. **(A)** Schematics of Experimental Design. Six to 8-week old male mice received subcutaneous (SC) injections of PCSK9 mutant AAV 2 weeks before the start of Western diet for 4 weeks and were infused with AngII (1,000 ng/kg/day, n=12-13 per group) for 28 days **(B)** Cumulative Incidence of Aortic Rupture. Thirty eight percent of WT and 0% of VAMP8^⁻/⁻^ died of aortic rupture before 28 days.**(C)** Representative Ultrasound Images. Proximal thoracic and abdominal aortas at week 4 (n=13 WT and n=9 VAMP8^⁻/⁻^). **(D)** Quantification of Aortic Diameter measured by ultrasonography. **(E)** Representative gross aortic images of study mice at termination. **(F)** Quantification of maximal abdominal aortic diameter n=13/group (control) and n=9 (AngII). Data are presented with box plots as mean ± SEM. The Kaplan-Meier survival log-rank test was used for survival curve analysis. The P-value for aortic diameter quantification was determined by Welch’s t-test. *** P ≤ 0.001. WT: wild type, AngII: angiotensin II, AoD: aortic diameter, Asc: ascending, Abd: abdominal

### VAMP8 Deficiency Alters Platelet and Aortic Transcriptomes

Given the protective phenotype observed in VAMP8^⁻/⁻^ mice, we sought to understand the underlying mechanisms, beyond a simple effect on secretion, by examining transcriptomic alterations in both platelets and aortic tissues from untreated animals to assess basal differences resulting from VAMP8 loss. Bulk RNA sequencing of VAMP8-deficient platelets versus WT revealed a distinct RNA profile, with 997 DEGs—520 upregulated and 477 downregulated **(Figure 6A)**. Reactome pathway analysis indicated significant enrichment in pathways related to platelet signaling and AAA, such as signaling by TGF-βeta family members **(Figure 6B)**. Notably, key genes critical for platelet function—including *Selp*, *Itga2*, *Gp6*, and *Ptgs1*—were significantly downregulated in VAMP8-deficient platelets **(Figure 6C)**. The aortic transcriptome of VAMP8^⁻/⁻^ mice showed significant downregulation of genes and pathways involved in platelet adhesion to exposed collagen and platelet activation **(Figure 6 D and E)**. This suggests that VAMP8 deficiency may reduce platelet-endothelial interactions and thrombotic potential within the aortic environment, aligning with the observed protection against AAA development and in addition to its effects on secretion, the loss of VAMP8 affects platelet biogenesis and alters their “program,”

**Figure 6:**
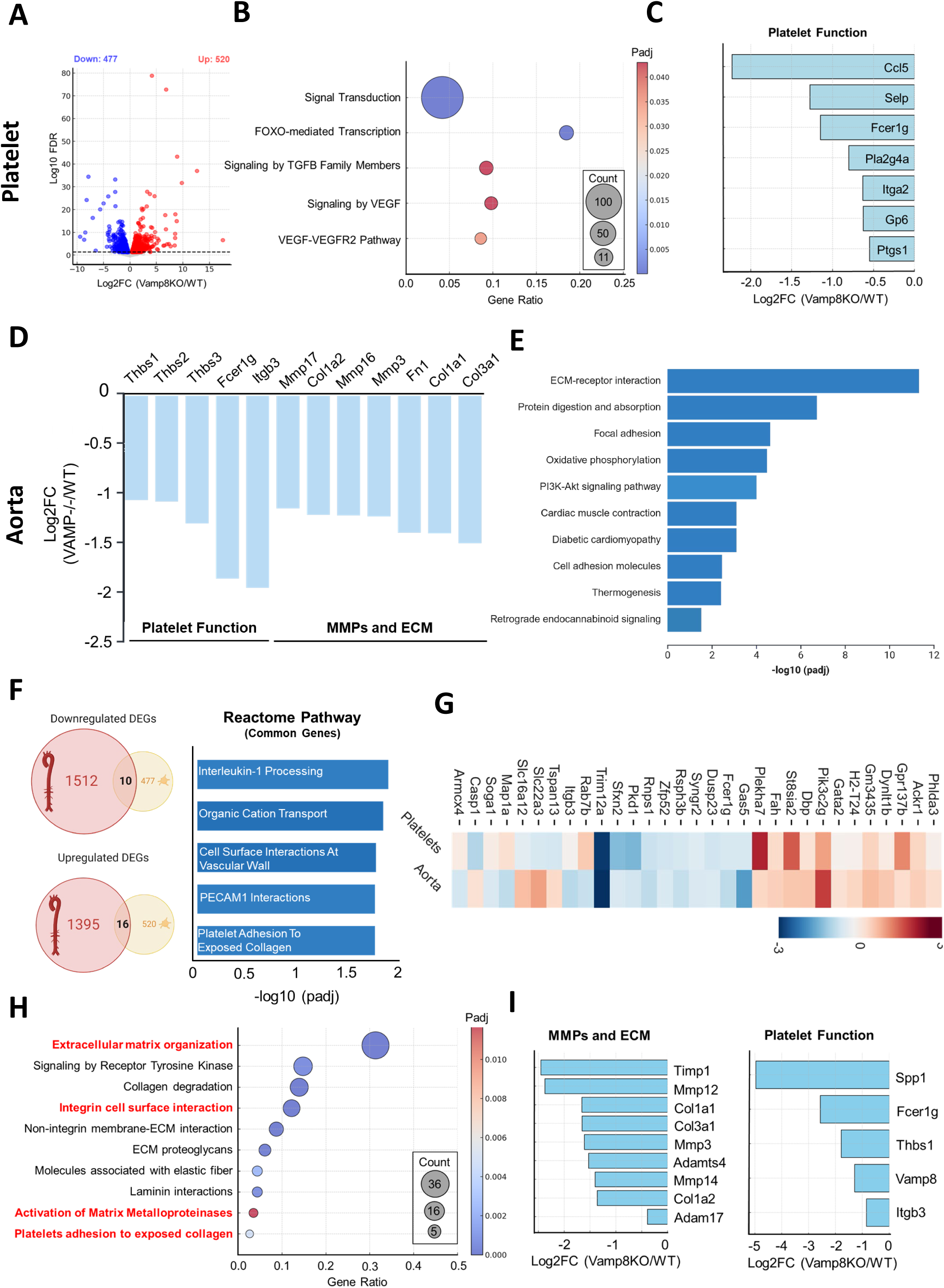
VAMP8 Deficiency Alters Platelet and Aortic Transcriptomes. **(A)** Volcano plot for differentially expressed genes between WT versus VAMP8 deficient platelets (n=2/group, each sample is consisted of pooled platelets from 2 mice). **(B)** The 5 most significant Reactome terms for platelet RNA sequencing data from WT vs. VAMP8 deficient mice. **(C)** Log fold change of genes related to platelet function compared between WT and VAMP8 deficient platelets. **(D)** Log fold change of genes related to matrix metalloproteinase (MMP) and extracellular matrix (ECM), chemokines and cytokines, and platelet function compared between WT and VAMP8 deficient aortic tissue. **(E)** Top 10 annotations of enrichment analyses in the Gene Ontology molecular function for WT versus VAMP8 deficient aortic tissue. **(F)** *Venn diagrams* and top annotations of Gene Ontology molecular function enrichment analysis for shared differentially expressed genes between downregulated and upregulated aortic and platelet datasets under basal condition between WT and VAMP8 deficient samples. **(G)** Heatmap with Z score coloring for common genes between platelet and aortic samples. **(H)** The 10 most significant Reactome terms and **(I)** Log fold change of genes related to MMP, ECM and platelet function compared between 5 day AngII infused aortic tissue of WT and VAMP8 deficient mice.

We examined common altered genes in both aortic and platelet transcriptomes in response to VAMP8 deficiency. This shared analysis revealed significant changes in key pathways that may collectively contribute to the vascular protective effects observed in VAMP8-deficient mice such as “interleukin-1 processing” and “organic cation transport,” highlighting a reduction in pro-inflammatory cytokine signaling and ion transport, which may decrease inflammatory stress within the vascular wall **(Figure 6F).** Additionally, enrichment of pathways related to “cell surface interactions at the vascular wall” and “PECAM1 interactions” suggests diminished leukocyte-platelet and endothelial interactions. Genes such as *Tspan13*, *Tap2*, *Syt2*, and *Rap2b* exhibited notable downregulation in both tissues, suggesting suppression of vesicle trafficking and signaling pathways that could potentially modulate platelet activation and vascular responses **(Figure 6G)**. Conversely, upregulated genes like *Steap2*, *Dbp*, and *Gpx3*, which may be involved in antioxidative responses and vascular repair mechanisms, were observed **(Figure 6G)**.

To further elucidate the role of VAMP8 in vascular responses, we assessed early transcriptional changes in the aorta after a short-term (5-day) AngII infusion in both VAMP8^⁻/⁻^ and WT mice. There was enrichment of pathways associated with “ECM organization,” “integrin cell surface interaction,” and “platelet adhesion to exposed collagen” (**Figure 6H**). The enrichment of these pathways suggests a blunted remodeling and adhesive response, which may mitigate the initial phases of vascular injury and inflammation induced by AngII. Furthermore, the downregulation of genes related to platelet function alongside the downregulation of MMPs indicates potential stabilization of the extracellular matrix in VAMP8^⁻/⁻^ mice **(Figure 6I)**.

## DISCUSSION

Our study provides robust evidence that VAMP8 deficiency offers significant protection against the development and progression of AngII-induced AAA. While platelets are not the only cells to express VAMP8, it is the primary v-SNARE in platelets and its deletion affects release from all three platelet granule types ^15,27,30^ and occlusive thrombosis **(Figure 4)**. Platelets are traditionally recognized for their roles in hemostasis and thrombosis. However, their contribution to the pathogenesis of AAA is garnering increasing attention ^7,8,31–34^. Our data indicate that platelet counts are significantly reduced in AngII-infused mice at both pre-overt pathology and pathological conditions, suggesting platelet consumption likely due to their activation by/within the aneurysmal sac. This observation aligns with previous findings indicating the presence of platelets in the ILT of human AAAs and highlighting their role in promoting inflammation and ECM degradation through interactions with macrophages and fibroblasts. ^8^ VAMP8 is essential for platelet granule secretion. The absence of VAMP8 especially impairs the release of α-granule contents, which include a myriad of bioactive molecules such as growth factors, chemokines, and cytokines ^18^. While VAMP8 deletion does not ablate hemostasis, it does affect occlusive thrombosis in a large caliber vessel (*i.e.,* carotid) suggesting that is important for platelet deposition and thrombus growth. Such a role could be important in the development of aneurysm where response to an initiating vascular insult or continued effects on damaged vessel walls could be mediated by platelet adherence, activation and secretion. ^15,18,27^ Our data showed that VAMP8^-/-^ mice show a significantly reduced aortic dilation and rupture rates in AngII-induced AAA models, suggesting that impaired platelet secretion confers a protective effect against AAA progression. These finding corroborates previous preclinical and clinical work that inhibiting platelets and modulating platelets activation in established AAA reduced thrombotic risk and progression of the pathology in AAA patients. ^3,7,35,36^

Our transcriptomic data following short-term AngII infusion **(Figure 3)** indicate that platelet function is altered in the early stages of disease development, which can be mitigated by VAMP8 deficiency. Transcriptome analyses of platelets from both mouse and human RNA sequencing data reveal an upregulation of genes and enrichment in pathways related to platelet activation and degranulation. This includes an increase in MMPs and enhanced ECM degradation, potentially accelerating the proteolytic breakdown of critical structural ECM components. Such degradation may heighten local inflammation and elicit a stronger aortic tissue response at the onset of pathology, contributing to compromised aortic wall structure. Further transcriptomic analysis of aortic tissue from short-term infused samples shows significant enrichment in pathways associated with ECM degradation and upregulation of genes linked to platelet function, inflammation, and MMPs. These factors—present within platelet cargo—are influenced by VAMP8 deficiency, suggesting a “platelet-aorta axis” mechanism that drives the initiation and progression of AAA. Our RNA sequencing analysis of suprarenal aortic tissues from VAMP8-deficient and WT mice also identified significant transcriptomic alterations at basal level. Specifically, VAMP8 deficiency was associated with the downregulation of genes encoding ECM-degrading enzymes and MMPs. These transcriptomic changes could contribute to the observed protection against AAA in VAMP8^-/-^ mice by enhancing aortic wall stability and reducing ECM turnover. Our data suggest that VAMP8 deficiency provides a more comprehensive protective effect by not only impairing platelet secretion but also altering the transcriptome to bolster aortic wall integrity. The clinical relevance of our findings is highlighted by the observation of reduced platelet counts and increased platelet activation in AAA patients, which is consistent with previous reports. ^2,7,8^ These findings suggest that platelets play crucial roles in AAA pathogenesis through their contribution to ILT formation and aortic wall degradation. While antiplatelet therapies have shown mixed results in clinical studies, our data highlight the potential of targeting specific components of the platelet secretory machinery, such as VAMP8, to prevent AAA progression.

Of note, transcriptomic changes in platelets were observed after just five days of AngII infusion, despite the platelet half-life being approximately four days. ^37^ This rapid alteration, including changes in VAMP8-deficient platelets, suggests potential upstream effects at the level of megakaryocytes (MKs)—the progenitor of circulating platelets. ^37,38^ These findings raise the possibility that AngII infusion and VAMP8 deficiency could influence thrombopoiesis at the megakaryocyte level, resulting in altered platelet production. The accompanying RNA-seq data imply that AngII infusion may also impact the way cargo is packaged in platelet granules early in disease progression. Such alterations could contribute to platelet-driven inflammatory and proteolytic responses observed in the disease model. Further studies are necessary to investigate these mechanisms and to clarify how changes in megakaryocyte activity and granule packaging might drive platelet function in the context of aortic pathology.

Our study provides novel insights into the role of platelet granule secretion in AAA development, complementing existing literature. Studies have shown that aspirin therapy, which modulates platelet function, improves fibrin clot characteristics and reduces thrombotic risk in AAA patients.^35^ In contrast, studies on ticagrelor underscores the complexity of platelet involvement in AAA, as it showed no significant effect on AAA growth despite effective platelet inhibition. ^6^. Others reported that antiplatelet therapy in patients with AAA but without symptomatic atherosclerosis showed limited effectiveness in reducing ischemic events and a trend towards higher bleeding risk, highlighting the need for careful consideration in prescribing such therapies. ^34^ Our data on VAMP8 deficiency suggest that targeting platelet secretion pathways could offer a more focused strategy for stabilizing the aortic wall and reducing inflammation. The protective effects of VAMP8 deficiency in AAA may be attributed to multiple mechanisms. Firstly, impaired platelet secretion reduces the release of pro-inflammatory and ECM-degrading molecules, thereby mitigating inflammation and ECM degradation in the aortic wall. Secondly, the transcriptomic changes observed in VAMP8-deficient mice indicate a shift towards enhanced ECM stability and reduced MMP activity, further contributing to aortic wall integrity. Future research should focus on developing targeted therapies that modulate VAMP8-mediated platelet secretion. Such interventions could provide a novel approach to preventing AAA progression by reducing platelet activation and granule release, thereby stabilizing the aortic wall and minimizing inflammation.

This study has several strengths. It is the first to demonstrate the protective effects of VAMP8 deficiency against AAA, providing new insights into the role of platelet granule secretion in AAA pathogenesis. The integration of transcriptomics data with physiological observations offers a comprehensive understanding of the molecular mechanisms underlying the protective effects of VAMP8 deficiency. Additionally, our findings are highly relevant to human AAA, as they align with clinical observations of platelet activation and function in AAA patients. However, there are also limitations to our study. The exclusive use of male mice may limit the generalizability of the findings to female populations, even though AAA is more prevalent in males.^39^ The reliance on a single animal model (AngII-infused mice) may not fully capture the complexity of AAA pathogenesis in humans, and additional models could provide a more comprehensive understanding. However, the AngII-induced AAA model closely replicates key aspects of human aneurysm pathology. This model effectively mirrors the characteristic luminal dilation, medial elastin fragmentation, atherosclerotic plaque formation, inflammatory cell infiltration, and proteolytic tissue degradation. Additionally, it also forms the extraluminal thrombus, providing a comprehensive representation of the disease process seen in human AAAs.^9,40^ Furthermore, our study used a global VAMP8^-/-^ model rather than a platelet-specific VAMP8 deficiency. This global knockout approach does not allow us to isolate the effects of VAMP8 deficiency specifically within platelets, as VAMP8 is also expressed in other cell types, including endothelial cells and smooth muscle cells, which might contribute to the observed protective effects. Future studies should employ cell-specific VAMP8 knockout models to delineate the exact role of platelet-derived VAMP8 in AAA pathogenesis.

## CONCLUSION

In conclusion, our study demonstrates that VAMP8 deficiency profoundly attenuates AngII-induced AAA by impairing platelet granule release and altering the transcriptomic profile of the aortic wall defining a potential “platelet-aorta axis” in disease initiation and progression. These findings highlight the critical role of platelet secretion in AAA pathogenesis and suggest that targeting VAMP8 could provide a novel therapeutic strategy for managing this life-threatening condition. Further research is warranted to translate these findings into clinical practice and explore the potential of VAMP8 inhibition as a treatment for AAA.

## ACKNOWLEDGMENTS

The authors wish to thank the members of the Whiteheart lab for their comments on the manuscript and their careful editing of the final version. S.M. designed the study, performed all experiments, acquired and analyzed the data, and wrote the manuscript. H.A. and ED assisted with experimental procedures. S.J contributed to the platelet transcriptome experiments. S.W.W. is the project leader, overseeing the entire project, and provided critical revisions to the manuscript. All authors reviewed and approved the final manuscript.

## SUPPLEMENTAL METHODS

The data supporting the findings of this study are available from the corresponding author upon reasonable request.

### Flow Cytometry Analysis

Washed mouse platelets (20 μL of 5 × 10^7^/mL) were analyzed either in a resting state (no agonist) or following stimulation with thrombin (0.05 U/mL) for 2 minutes at room temperature. The reactions were halted by adding a two-fold excess of hirudin. Platelets were then incubated with 2.5 μL of FITC-conjugated antibodiy for 20 minutes at 37° C. Subsequently, the platelets were diluted 10-fold with HEPES-Tyrode’s buffer (pH 6.5) and transferred to polystyrene FalconTM tubes (BD Biosciences, San Jose, CA). Fluorescence intensity was measured using a BD FACSymphonyTM flow cytometer (BD Biosciences, San Jose, CA). Platelet populations were identified by adjusting the voltages for forward scatter (FSC) and side scatter (SSC). Fluorescence intensity was optimized by adjusting the voltages for the green (FITC) and yellow (PE) excitation channels. Data analysis was performed using FlowJoTM software v10.8.0 (BD Biosciences, San Jose, CA). For each sample, 50,000 events were recorded, and geometric mean fluorescence intensities (GFMI) were calculated and plotted with statistical analysis.

### Systolic Blood Pressure Measurements

Systolic blood pressure was assessed using a non-invasive tail cuff system (Coda 8, Kent Scientific), following the protocol described previously ^41^. Conscious mice were placed in restrainers on a heated platform. Blood pressure readings were taken 20 times daily at the same time for three consecutive days. Data points with values <60 or >250 mmHg, standard deviation >30 mmHg, or fewer than 5 valid cycles out of 20 were excluded from analysis.

### Aortic Tissue Processing, Histological Analyses, and Immunostaining

Aortic tissue was harvested after a 28-day period for histological analyses. Mice were euthanized by CO_2_ inhalation, followed by opening the thoracic cavity and perfusing saline (10 ml) through the left ventricle. For immunostaining, suprarenal aortas were dissected free and placed in OCT compound for sectioning. Serial cross-sections (10 μm) were collected as frozen sections and incubated with acetone for 10 minutes at −20°C. Subsequently, the sections were incubated with goat serum for 1 hour at 4°C. Cryosections were then incubated with a blocking solution (5% goat serum, 0.5% Triton X-100 in PBS) for 1 hour. The sections were incubated with primary antibodies diluted in the blocking solution overnight at 4°C. The primary antibodies used were Rabbit monoclonal Recombinant Alexa Fluor® 488 Anti-VAMP8/EDB antibody [EP2629Y] (ab202828) and Recombinant Alexa Fluor® 568 Anti-CD42b antibody [SP219] (ab312777). The following day, sections were washed with 10X ImmunoStain wash buffer (GeneTex GTX73344) and cover slipped. DAPI (Sigma-Aldrich, D9542) was used for nuclear staining. Hematoxylin and eosin staining (catalog No. 26043-06; Electron Microscopy Sciences; catalog No. AB246824; abcam) and Masson’s trichrome staining (following the manufacturer’s protocol, Polysciences, Warrington, PA, USA) were also performed. Images of histological and immunostaining were captured using an Axioscan Z7 (Zeiss) or Nikon AXR Inverted Confocal Microscope.

### Measurements of abdominal aortic diameters by Ultrasonography

The ascending or abdominal aorta was imaged in vivo using a Vevo 3100 ultrasound system with a 40-MHz transducer as described previously ^42^. Mice were anesthetized with isoflurane (1.0-2.5% vol/vol), maintaining a heart rate between 400-550 beats per minute during imaging. Aortic luminal diameter measurements were taken from inner edge to inner edge at end diastole, averaged over three distinct heartbeats. An independent investigator, blinded to the study groups, verified all measurements.

### *Ex Vivo* Aortic Diameter Measurement

Mice that survived until the end of the study were euthanized by CO₂ inhalation. Following euthanasia, the right atrial appendage was excised, and approximately 10 mL of saline was perfused through the left ventricle to flush the circulatory system. The aortic tissue was then carefully harvested, and fixed in neutrally buffered formalin (10% vol/vol) overnight at room temperature. Periaortic adventitia were removed carefully after the fixation. Subsequently, the aortas were pinned on a black wax surface. Images of the aorta were captured using a Nikon SMZ stereoscope (models 25 or 800; Nikon Corporation). The captured images were analyzed using NIS-Elements AR software (version 5.11; Nikon Corporation). To measure the aortic diameters, a measurement line was drawn perpendicularly to the aortic axis at the most dilated area of the suprarenal aortic region. The measurement software was calibrated for each image using the ruler included in the image to ensure accurate scaling.

### *En Face* Quantification of Atherosclerosis

An *en face* method, in accordance with American Heart Association guidelines ^43^, was used to measure atherosclerotic lesions. The intimal surface of the aorta was exposed through longitudinal cuts and pinned onto a black wax surface. *En face* images were captured using a Nikon SMZ stereoscope (models 25 or 800; Nikon Corporation) equipped with a digital camera. Lesions were manually traced from the ascending aorta to the descending thoracic aorta, extending 1 mm distal from the left subclavian artery. The captured images were analyzed using NIS-Elements AR software (version 5.11; Nikon Corporation). The extent of atherosclerosis was quantified by calculating the percentage of the lesion area relative to the total intimal area of the examined aortic region.

### Total Cholesterol Measurement

Blood samples were collected from mice at termination via the right ventricle using EDTA (1.8 mg/mL). The collected blood was then centrifuged at 3,000 X g for 10 min at 4°C to obtain plasma for analysis. Mouse plasma lipid concentrations were analyzed using an enzymatic kit (Cat. No. C7510-120; Pointe Scientific).

## MAJOR RESOURCES TABLES

### Animals (in vivo studies)

| Mouse Strain | Sex | Strain # | Persistent ID/URL |
| --- | --- | --- | --- |
| C57BL/6J | Male | 000664 | https://www.jax.org/strain/000664 |
| VAMP8<sup>-/-</sup> | Male | *In-house* | N/A |

### Primary Antibodies for Immunostaining

| Antibody | Vendor | Cat# | Working Concentration |
| --- | --- | --- | --- |
| Alexa Fluor® 488 Anti-VAMP8/EDB | Abcam | ab202828 | 3 µg/mL |
| Alexa Fluor® 568 Anti-CD42b | Abcam | ab312777 | 3 µg/mL |
| Rabbit nonimmune IgG | ImmunoReagents | Rb-003-V | 3 µg/mL |

### Animal Study Information Following the ARRIVE Essential 10

| Figure | Groups | Sex | Age (Weeks) | Number (Start) | Number (Termination) |
| --- | --- | --- | --- | --- | --- |
| **1 A** | Control | Male | 12 | 8 | 8 |
| **1 A** | AngII | Male | 12 | 8 | 8 |
| **1 B** | Control | Male | 12 | 13 | 9 |
| **1 B** | AngII | Male | 12 | 13 | 9 |
| **2 A-C** | Control | Male | 8 | 4 | 4 |
| **2 A-C** | AngII | Male | 8 | 4 | 4 |
| **2 G-J** | Control | Male | 8 | 3 | 3 |
| **2 G-J** | AngII | Male | 8 | 3 | 3 |
| **3 A-D** | Control | Male | 8 | 4 | 4 |
| **3 A-D** | AngII | Male | 8 | 4 | 4 |
| **4 A-C** | WT | Male | 8-12 | 5 | 5 |
| **4 A-C** | VAMP8<sup>-/-</sup> | Male | 8-12 | 5 | 5 |
| **5 B-F** | WT | Male | 8-12 | 12 | 9 |
| **5 B-F** | VAMP8<sup>-/-</sup> | Male | 8-12 | 13 | 13 |
| **6 A-I** | Control | Male | 8 | 4 | 4 |
| **6 A-I** | AngII | Male | 8 | 4 | 4 |

## ARRIVE ESSENTIAL 10 CHECKLIST

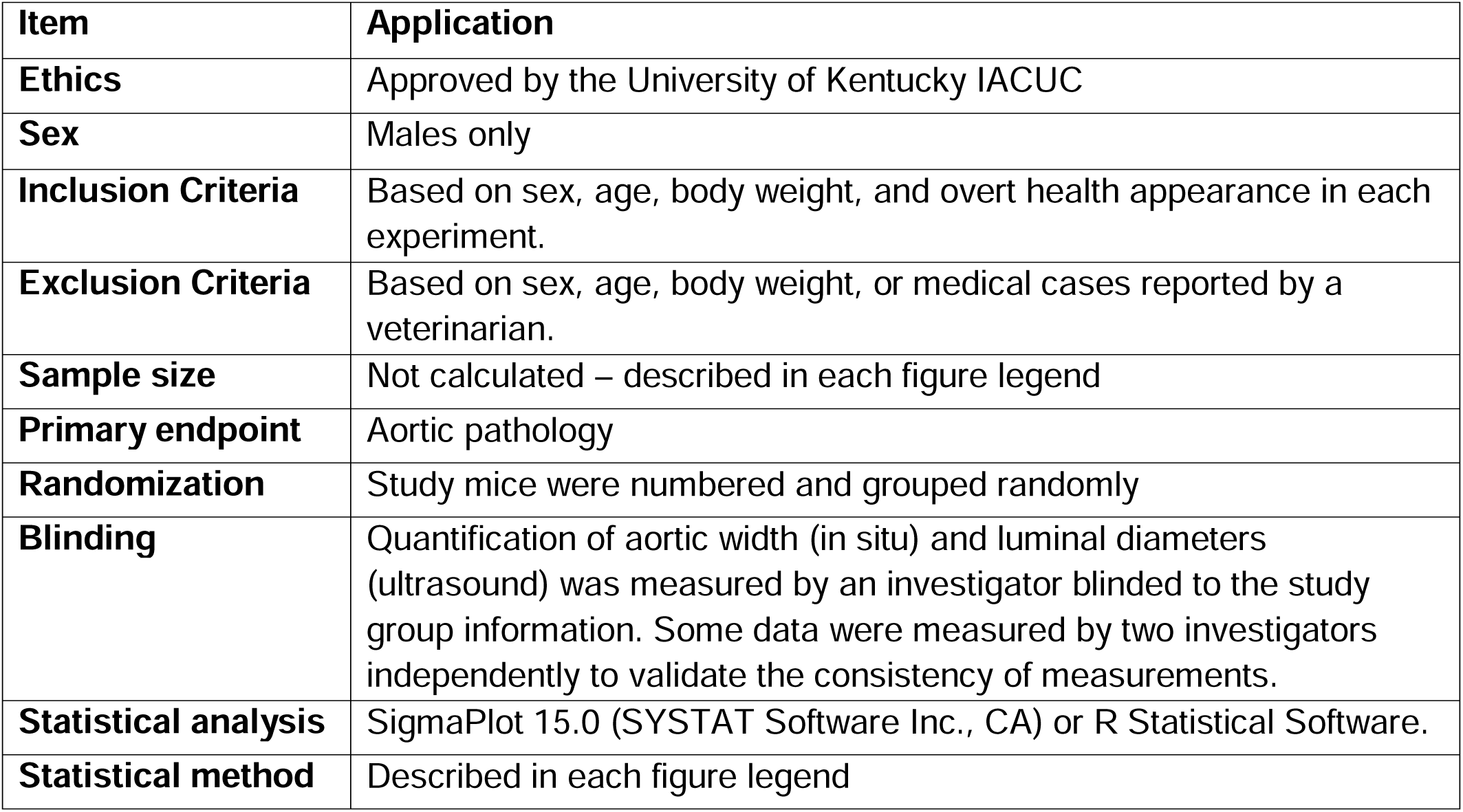

| **Item** | **Application** |
| --- | --- |
| **Ethics** | Approved by the University of Kentucky IACUC |
| **Sex** | Males only |
| **Inclusion Criteria** | Based on sex, age, body weight, and overt health appearance in each experiment. |
| **Exclusion Criteria** | Based on sex, age, body weight, or medical cases reported by a veterinarian. |
| **Sample size** | Not calculated – described in each figure legend |
| **Primary endpoint** | Aortic pathology |
| **Randomization** | Study mice were numbered and grouped randomly |
| **Blinding** | Quantification of aortic width (in situ) and luminal diameters (ultrasound) was measured by an investigator blinded to the study group information. Some data were measured by two investigators independently to validate the consistency of measurements. |
| **Statistical analysis** | SigmaPlot 15.0 (SYSTAT Software Inc., CA) or R Statistical Software. |
| **Statistical method** | Described in each figure legend |

**Supplemental Table II.** Mouse housing conditions.

|  | Mouse Housing Condition | Note |
| --- | --- | --- |
| Set temperature range | 68-74°F (20-23°C) |  |
| Set humidity | 50% |  |
| Light cycle (light:dark) | 14:10 hours |  |
| Water | Reverse osmosis water | ad libitum |
| Feed | 18% Protein Rodent Diet, #2918,<br>Teklad Irradiated Global (Envigo) | ad libitum |
| Bedding | Aspen hardwood chips, #7090A,<br>Harlan Teklad Global (Envigo) |  |
| SPF (Specific pathogen free) | Yes |  |

## Supplementary Figures

**Supplementary Figure 1:**
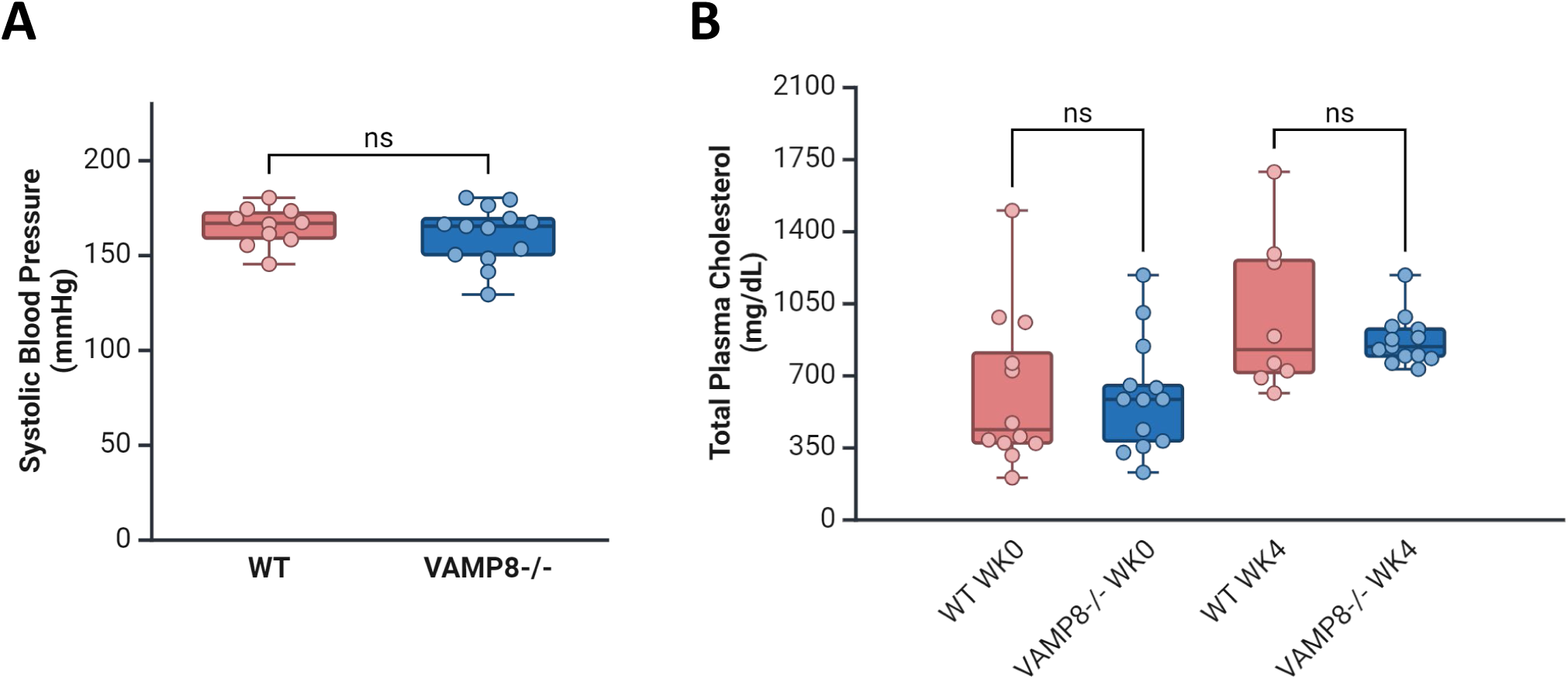
VAMP8 Deficiency Does Not Affect Total Cholesterol or Blood Pressure. **(A)** Systolic blood pressure in study mice measured by the tail-cuff method. Blood pressure was measured 20 times at the same time each day for three consecutive days. **(B)** Total plasma cholesterol at week 0 (2 weeks post-PCSK9 mutant AAV injection) and week 4 (6 weeks after PCSK9 injection). Data are presented with circles and box plots as mean ± SEM. The P-value for blood pressure was determined by Student’s t-test. For total cholesterol, the P-value was calculated by two-way analysis of variance followed by the Holm-Sidak test. N.s; not significant.

**Supplementary Figure 2:**
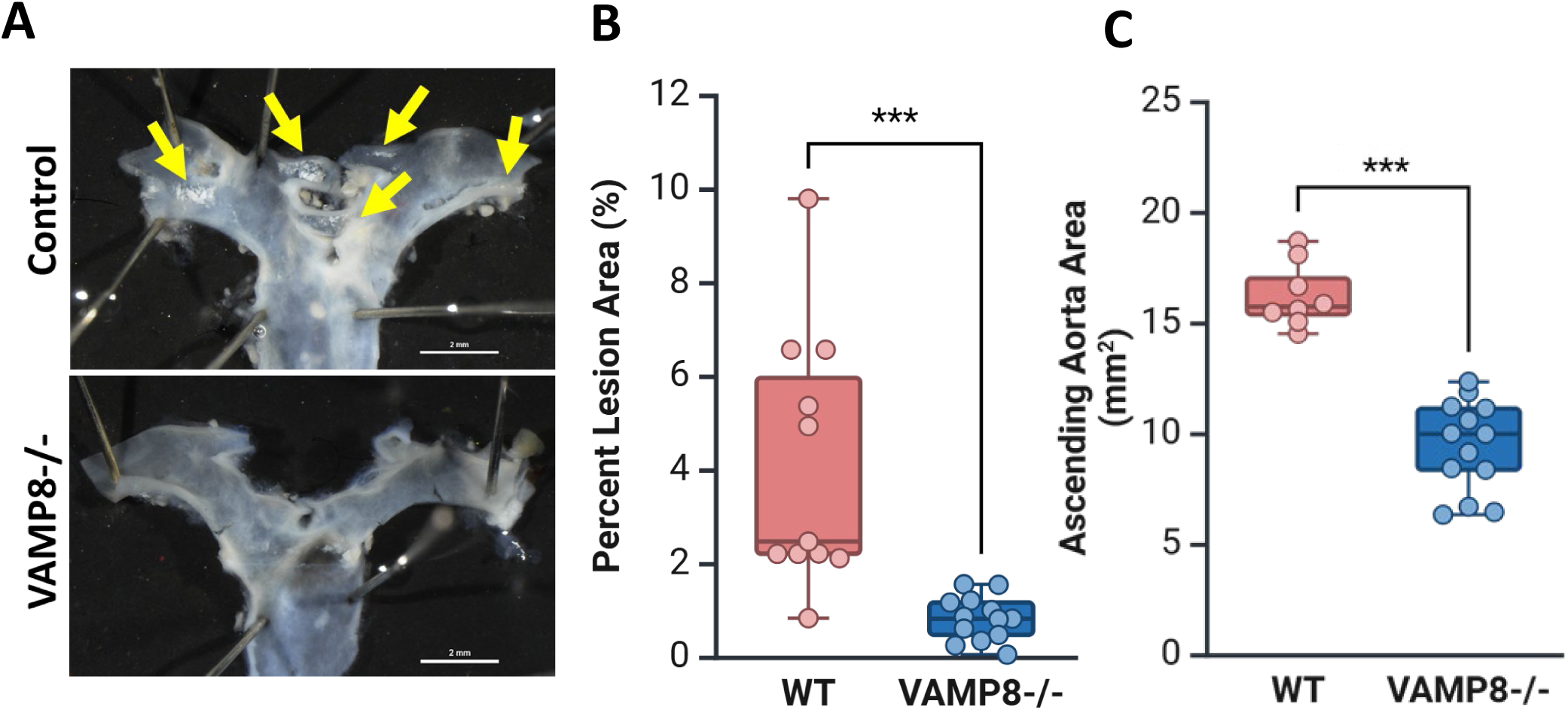
VAMP8 Deficiency Suppressed Early-stage Atherosclerosis Development. **(A)** Representative gross images of atherosclerosis formation in study mice (n=9 WT, n=12 VAMP8-/-) **(B)** Percent atherosclerotic lesion area and **(C)** ascending aortic area quantified by an *en face* approach. Yellow arrows indicate atherosclerotic lesion areas. Data are presented with circles and box plots as mean ± SEM. P-value was determined by Mann-Whitney U test. *** P ≤ 0.001

